# Vaccine Hyporesponse Induced By Individual Antibiotic Treatment In Mice And Non-Human Primates Is Diminished Upon Recovery Of The Gut Microbiome

**DOI:** 10.1101/2021.04.02.438241

**Authors:** Gokul Swaminathan, Michael Citron, Jianying Xiao, James E Norton, Abigail L Reens, Begüm D. Topçuoğlu, Julia M Maritz, Keun-Joong Lee, Daniel C Freed, Teresa M Weber, Cory H White, Mahika Kadam, Erin Spofford, Erin Bryant-Hall, Gino Salituro, Sushma Kommineni, Xue Liang, Olga Danilchanka, Jane A Fontenot, Christopher H Woelk, Dario A Gutierrez, Daria J Hazuda, Geoffrey D Hannigan

## Abstract

Emerging evidence demonstrates a connection between microbiome composition and suboptimal response to vaccines (vaccine hyporesponse). Harnessing the interaction between microbes and the immune system could provide novel therapeutic strategies for improving vaccine response. Currently we do not fully understand the mechanisms and dynamics by which the microbiome influences vaccine response. Using both mouse and non-human primate models, we report that short-term oral treatment with a single antibiotic (vancomycin) results in disruption of the gut microbiome and this correlates with a decrease in systemic levels of antigen-specific IgG upon subsequent parenteral vaccination. We further show that recovery of microbial diversity before vaccination prevents antibiotic-induced vaccine hyporesponse, and that the antigen specific IgG response correlates with the recovery of microbiome diversity. RNA-sequencing analysis of small intestine, spleen, whole blood, and secondary lymphoid organs from antibiotic treated mice revealed a dramatic impact on the immune system, and a muted inflammatory signature is correlated with loss of bacteria from *Lachnospiraceae, Ruminococcaceae*, and *Clostridiaceae*. These results suggest that microbially modulated immune pathways may be leveraged to promote vaccine response and will inform future vaccine design and development strategies.

**Importance:** Antibiotic-induced gut microbiome disruption has been linked to reduced vaccine efficacy. Despite these observations, there remains a knowledge gap in the specific mechanisms by which antibiotics and the gut microbiome influence vaccine efficacy. We aim to contribute to the field’s growing mechanistic understanding by presenting a detailed analysis of antibiotic treatment and recovery as it relates to vaccine response and the microbiome. Using animal models, we show that short-term antibiotic treatment prior to vaccination results in diminished vaccine-specific immune responses, and that these are correlated with specific microbiome signatures. We also demonstrate the converse, in which gut microbiome recovery can result in improved vaccine response. We further reveal that antibiotics can significantly alter multiple relevant immune pathways and this alteration in immune tone may contribute to the vaccine hyporesponse. We expect our findings will enable the continued prosecution of the role of the microbiome in modulating the host immune system.

## Introduction

Vaccination plays an essential role in global health by contributing to prevention and eradication of infectious diseases. When individuals are immunized with vaccine antigens, the body mounts an immune response that leads to long-term immune memory and protection against future infection. However, many vaccines still suffer from relatively low response rates in some populations or geographical locations (1–5). The key immunological mechanisms underlying this lack of response are not well understood, which limits progress in improving vaccine outcomes. Investigating the pathways involved in such vaccine immune hyporesponsiveness may inform new solutions to increasing vaccine efficacy and facilitate greater protection in at-risk populations (6, 7).

Emerging evidence in clinical and preclinical studies suggests that vaccine hyporesponse may be linked to changes in the human microbiome — the community of microbes that live in and on the human body (8). Found in areas such as the gut, the skin, and respiratory and genitourinary mucosal surfaces, the microbiota are a rich source of nutrients, biologically active metabolites, and immunoregulatory molecules (9, 10). The microbiome plays a key role in numerous aspects of human biology (11–13), and perturbation of the gut microbiome has a significant influence on various disease outcomes (14–16). In particular, natural or antibiotic-induced dysbiosis of the gut microbiome has been associated with vaccine hyporesponse in humans (2, 17–21). Preclinical studies in antibiotic-treated or germ-free mice also support a connection between the microbiota and vaccine response (22–25).

Although multiple lines of evidence demonstrate a link between the microbiome and response to a variety of vaccines, we do not fully understand the underlying microbial and immunological mechanisms that dictate the breadth and longevity of vaccine-induced antigen-specific immune responses, nor a direct influence of the microbiome on such responses. Studies have suggested some correlation between microbiome and vaccine responses (e.g. inflammation and microbial secondary bile acids) (2, 22, 25–28), but they have yet to focus on microbiome perturbation with recovery dynamics in mice or non-human primates (NHPs). Here we begin to address this knowledge gap by characterizing the dynamics of vaccine hyporesponse associated with antibiotic-mediated microbiome perturbation and recovery in mice as well as non-human primates.

We report that short-term oral treatment of individual antibiotics prior to parenteral vaccination impacted antigen-specific immune responses, and that the extent of vaccine hyporesponse correlated with the extent of microbiome disruption across different antibiotics. Specifically, we found that vancomycin, a glycopeptide antibiotic that specifically targets gram-positive bacteria, robustly impacted vaccine-induced antigen-specific IgG responses. Furthermore, we observed that vancomycin treatment diminished IgG responses against multiple vaccines including an adjuvanted model sub-unit vaccine antigen (i.e., ovalbumin), as well two clinically relevant vaccines (i.e., hepatitis B virus vaccine and influenza vaccine). We report data to suggest that natural recovery of microbiome diversity over time after antibiotic treatment was sufficient to mitigate vancomycin-induced vaccine hyporesponse in both murine and non-human primates. We further identified immune signatures associated with potential commensal bacteria belonging to the taxa *Lachnospiraceae, Ruminococcaceae*, and *Clostridiaceae*. These results further support the link between the microbiome and immune system and also provide an improved understanding of the dynamic connection between recovery of the microbiome and vaccine response. Such an improved understanding of host and microbiome-derived factors is critical for optimal outcomes after vaccination and has the potential to influence the future design of vaccines and immunotherapies.

## Results

### Microbiome Perturbation by Different Antibiotics Differentially Modulates Vaccine Immune Response

Recent studies have demonstrated that treatment with a cocktail of antibiotics prior to vaccination alter vaccine response by disrupting the microbiome (2, 22, 24, 27–29). However, the majority of these studies utilized chronic treatment (~4 weeks) with a combination of antibiotics. We therefore hypothesized that short-term (i.e., 7 days) oral treatment of individual antibiotics from different classes may differentially impact vaccine response, due to differences in their impact on the microbiome. To test this hypothesis, we treated BALB/c mice for 7 days with each of seven antibiotics in drinking water with no artificial sweeteners or additives, followed by one day of recovery without antibiotics, prior to the first dose of intramuscular vaccination with alum-adjuvanted ovalbumin (OVA) antigen (**Figure 1A**). A booster dose with no preceding antibiotic treatment was delivered 30 days after the first dose (**Figure 1A**). To ensure that the antibiotics did not adversely affect the health of mice during treatment, we confirmed acceptable and consistent water intake, food intake, estimated dosage, and body weight throughout the antibiotic treatment period (**Figure S1**). Twelve days following the booster dose, we found that each antibiotic had a unique reductive impact on serum OVA-specific IgG titer compared to water-treated control mice (**Figure 1B**).

**Figure 1.**
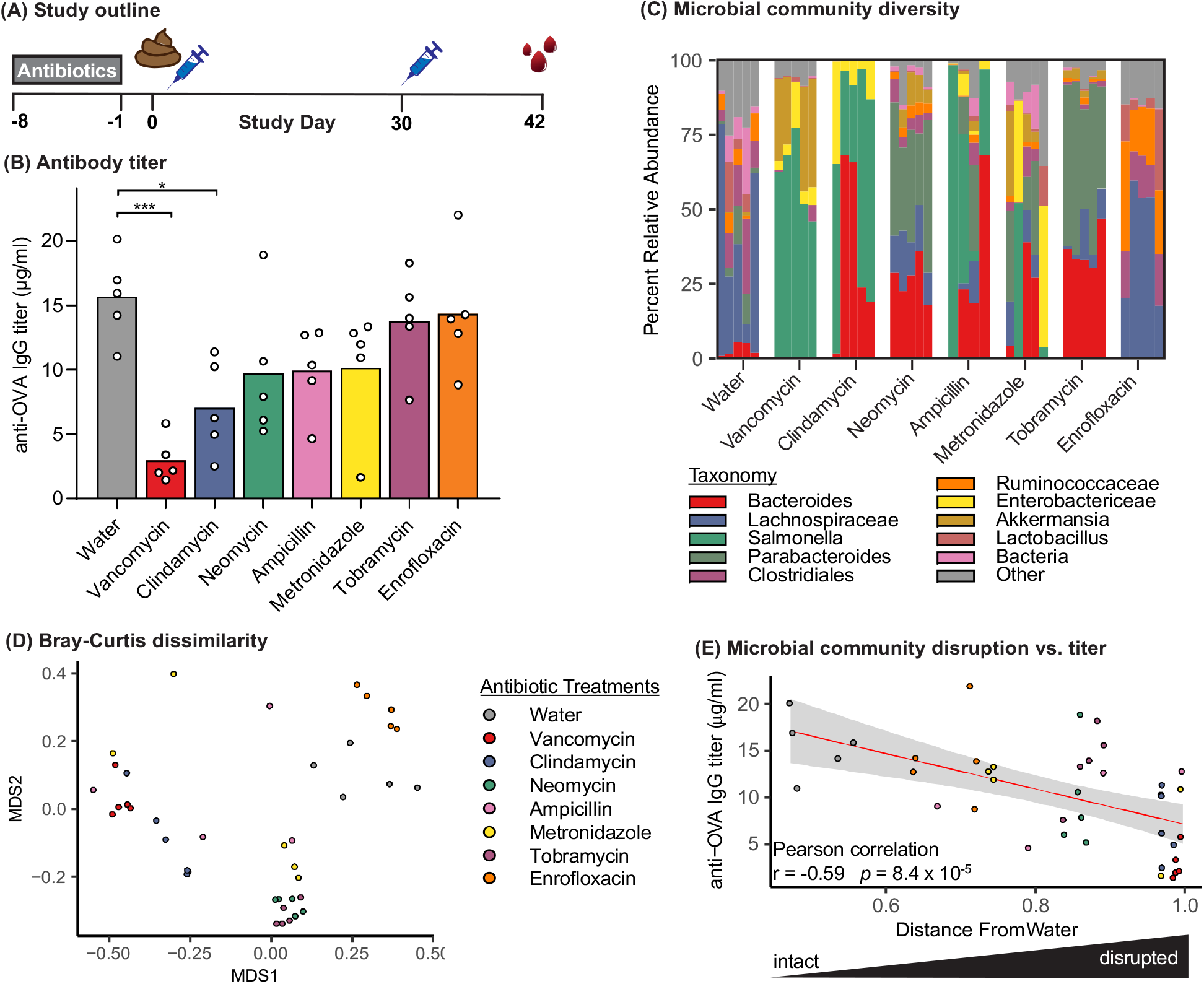
Oral administration of antibiotics differentially affects model sub-unit vaccine outcome and gut microbiome profiles. (A) Study design and dosing schedule. BALB/c mice were orally treated with antibiotics for seven days (gray rectangle), followed by a single day of no antibiotic treatment before a prime-boost OVA immunization regimen (blue syringes). Stool collection and harvesting of serum are indicated by brown stool and red icons, respectively. (B) Serum OVA-specific IgG titers at Day 42 after immunization. Measurements from individual mice (circles) are superimposed on group mean (bar). * p < 0.05, *** p <0.001 compared to water by one-way ANOVA with Dunnett’s post-hoc test. (C) Stacked bar plot of top 10 bacterial genera based on relative abundance determined by 16S rRNA gene sequencing of fecal samples collected at study day 0. Each column represents an individual mouse. (D) Non-metric multidimensional scaling (MDS) ordination of mouse microbiome communities with community dissimilarity measured using the Bray-Curtis algorithm. Each point represents the microbiome of an individual mouse; greater distances between points indicate less similar community composition. (E) Correlation scatter plot of OVA-specific IgG titer for each individual mouse and distance between the microbial community composition and the average composition of water-treated animals. Red line indicates linear regression line of best fit; gray shading represents 95% confidence interval.

We validated the pharmacodynamic impact of oral antibiotic treatment by characterizing the gut microbiome of each mouse at time of vaccination using 16S rRNA gene sequencing of fecal pellets. As expected, each antibiotic treatment uniquely altered the microbiome, which resulted in distinct microbial communities at the time of vaccination (**Figure 1C**). Microbial communities from mice treated with vancomycin and clindamycin, the two antibiotics with the strongest impact on vaccine response, visibly clustered together because of their shared community composition (**Figure 1D**), suggesting the presence of some common community elements in these most disruptive antibiotic treatments (e.g. Salmonella and Enterobactericeae).

Because we observed some community structure commonalities between our treatments, we performed a correlation analysis across all mice to determine whether specific community signatures were associated with resulting titer. We observed no strong correlations between the relative abundance of individual bacterial taxa and antibody titers (**Figure S2A**). We also investigated whether reduced antibody titer was associated with total community disruption (beta diversity compared to water) and observed an overall correlation between antibody titers and the degree of antibiotic-mediated microbiome disruption (**Figure 1E**). Together, these results suggest that the extent of microbiome community disruption at the time of vaccination correlated with the extent of vaccine hyporesponse and that this phenomenon may have been driven by changes in complex community interactions rather than by disruption of specific individual bacterial taxa.

### Antibiotic-mediated Vaccine Hyporesponse is Associated with Altered Microbiome Functionality

The correlation between antibody titers and bacteria at the community scale, rather than the individual genus level, suggested this complex dynamic could be informed by a function-based analysis. Microbes from different taxonomic groups can execute redundant functions and may sometimes be interchangeable (30), which could mask common functional alterations across antibiotic treatments in our data. We therefore supplemented our taxonomic analyses with shotgun metagenomic sequencing and identified correlations between the enrichment of microbiome functional potential and antibody titers (**Figure S2B-E**). The four pathways that significantly correlated with antigen specific titers were metabolic pathways such as terpenoid-quinone biosynthesis, sulfur metabolism, tryptophan metabolism, and selenocompound metabolism. All of the observed significant correlations were negative, meaning that enrichment of the metabolic pathways was associated with reduced antigen-specific IgG titers. The role of these microbial metabolic pathways in immune function and vaccine response is unclear, though they have previously been associated with microbial dysbiosis (31–34) and may result from expansion of normally rare microbes in antibiotic-treated mice (35).

In addition to correlating sample pathway enrichment with resulting vaccine titer, we also identified the metagenomic pathways depleted and enriched in each antibiotic treatment group relative to water treated controls (**Tables S1, S2, FDR corrected p-values < 0.01**). Four pathways were depleted in the antibiotic treatment groups, and while three of the pathways were only depleted in a single antibiotic group, the aminoacyl-tRNA biosynthesis pathway was depleted in five antibiotic treatment groups including clindamycin, metronidazole, neomycin, tobramycin, and vancomycin. This signal could reflect depletion of *Ruminococceae* and *Lactobacillales* across those antibiotics (36, 37). Conversely, there was no overlap of enriched pathways across antibiotic treatments, though many were associated with core metabolism and growth. The majority of enriched pathways (12 of 15) were in the vancomycin group, including lipopolysaccharide biosynthesis, which mirrors the enrichment of Gram-negative genera after vancomycin treatment (**Figure 1B**). Together, these data likely reflect alterations in representation of specific pathways due to reduced bacterial diversity after antibiotic treatment, especially in the vancomycin group, but future investigation may reveal functional links between altered microbial biosynthetic activity and vaccine response.

### Vancomycin-Induced Vaccine Hyporesponse is Minimized by Allowing Sufficient Time for the Microbiome to Recover

Our data above suggest that microbiome disruption at the time of immunization was associated with suboptimal antigen-specific IgG responses. To further test association of the microbiome with vaccine specific IgG titers, we asked whether the converse relationship was true: Can vaccine response be improved by allowing the disrupted microbiome to recover? We hypothesized that extending the time after antibiotic treatment and before immunization would allow for microbiome recovery and prevent vaccine hyporesponse in antibiotic-treated animals.

We tested our hypothesis using vancomycin as a model antibiotic due to its robust effect on humoral vaccine responses observed above (**Figure 1**). Vancomycin also demonstrated consistent vaccine-modulatory effects of subsequent studies wherein: i) vancomycin, but not ampicillin, robustly impacted alum-adjuvant OVA vaccine-specific titers in a different mouse strain (**C57Bl/6; Figure S3A**); ii) vancomycin decreased antigen-specific IgG titers and T cell responses when mice were vaccinated with OVA adjuvanted with a novel lipid nanoparticle (LNP)-based adjuvant (**Figure S4A-D**); and iii) vancomycin reduced antigen-specific IgG titers against a clinically-relevant hepatitis B vaccine conducted in two geographically distinct vivaria (**Figure S4E-F**). Vancomycin treatment has previously been shown to disrupt response to nonadjuvanted inactivated influenza vaccine (22) and rabies vaccine (38), and is included in many antibiotic cocktails used to perturb the microbiome for vaccination studies with variable effects on individual vaccine antigens (22, 27, 28, 38). Vancomycin treatment has also been shown to modulate several immune-related pathways (39–42), and reduces the abundance of bacteria belonging to *Clostridiales* and *Bacteroidales*, while allowing increases in *Proteobacteria* (39–43), in agreement with our data (**Figure 1C**).

To test our hypothesis using vancomycin, we treated mice with vancomycin and extended the number of days between antibiotic cessation and immunization with LNP adjuvanted-OVA vaccine from 1 to 7, 14, or 21 days (**Figure 2A**). We observed a visible trend between the length of recovery time after vancomycin treatment and antigen-specific antibody titers, in which animals with shorter recovery times had statistically significantly decreased antibody titers, whereas mice given 21 days of recovery before immunization had similar antibody titers to water-treated animals (**Figure 2B**).

**Figure 2.**
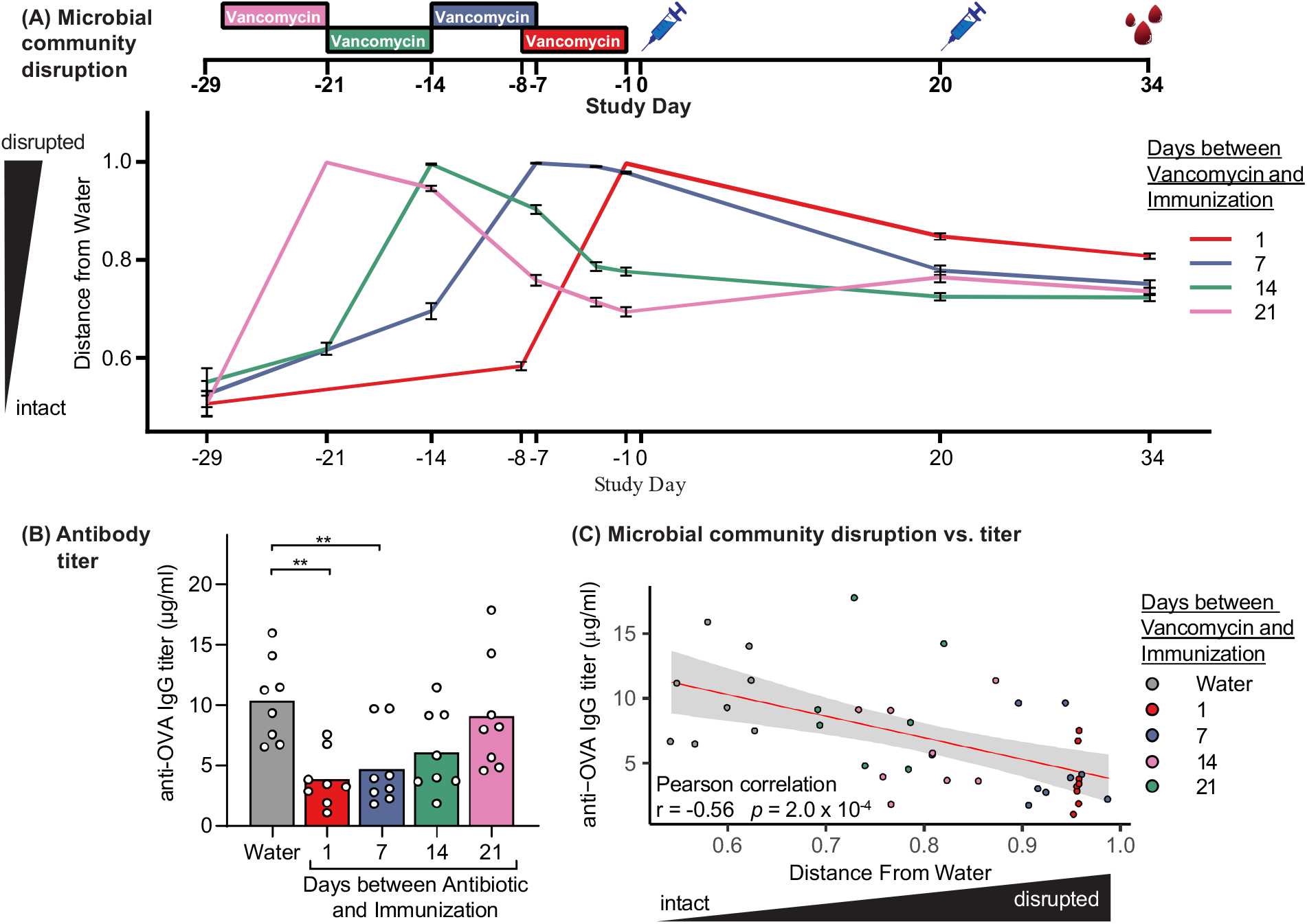
Extended interval between antibiotic treatment and immunization prevents antibiotic-induced vaccine hyporesponse. (A) BALB/c mice were orally treated with vancomycin for 7 days (colored rectangles), followed by 1, 7, 14, or 21 days without antibiotic treatment prior to immunization with OVA and VA-510 LNPs on Days 0 and 20 (blue syringes). Harvesting of serum is indicated by red droplets. Fecal samples collected the antibiotic and vaccination phases were subjected to 16s rRNA gene sequencing. The mean microbiome community dissimilarity compared to the average baseline community composition across all groups, calculated using the Bray-Curtis metric and plotted over time. Data are mean and standard error for each treatment group. (B) Serum anti-OVA IgG titers were measured at Day 34; measurements from individual mice (circles) are superimposed on group mean (bar). ** p < 0.01 compared to water by one-way ANOVA with Dunnett’s post-hoc tast. (C) Correlation scatter plot of each individual mouse’s OVA-specific IgG titer versus distance between the microbial community composition and the average composition of water-treated animals, measured at Day -1. Red line indicates linear regression line of best fit; gray shading represents 95% confidence interval.

Microbiome profiling by 16S rRNA gene sequencing revealed that microbial community recovery after vancomycin treatment followed defined dynamics, as measured by community dissimilarity (beta-diversity) from water (**Figure 2A**) and OTU richness (**Figure S5A**). Prior to vaccination, each treatment group had recovered to a different extent, and the extent of microbiome disruption was significantly negatively correlated with resulting antigen-specific antibody titers (**Figure 2C**). Of note, this result mirrors the correlation observed between antibody titers and microbiome disruption mediated by different classes of individual antibiotics (**Figure 1E**), demonstrating that microbiome disruption is associated with vaccine hyporesponse in both disruption and recovery models.

To confirm that these results are not due to differences in vancomycin treatment conditions between groups or direct effects of vancomycin on the systemic immune system, we validated that vancomycin exposure was comparable across mice, that >98% of vancomycin was excreted in the feces by the time of immunization relative to peak levels, and that vancomycin remained undetectable systemically in the blood at any timepoint (**Figure S5E,F**).

Because we found that greater disruptions of the microbiome were associated with greater reductions in the antigen-specific IgG response (**Figure 1E**), we hypothesized that reducing microbiome disruption by decreasing vancomycin treatment time to one or three days would also result in a reduced impact on vaccine hyporesponse. Indeed, we found that three days of vancomycin treatment moderately impacted the antigen-specific IgG titers following vaccination, while one day of treatment did not alter response (**Figure S5C**). We also confirmed that a single day of vancomycin treatment only moderately reduced bacterial richness compared to three or seven days of treatment (**Figure S5B**). The findings support our previously observed trends toward reduced vancomycin exposure in the gut (**Figure S5E,F**) and that the extent of microbiome disruption correlated with the effect on antigen-specific antibody response (**Figure 1**). Together, these findings suggest that mitigating the effects of antibiotic treatment on the gut microbiome by delaying immunization or decreasing treatment duration could prevent antibiotic-induced vaccine hyporesponse.

We further explored whether the reduction in antigen-specific IgG titers were correlated with individual bacterial taxa prior to immunization. We found no taxa strongly associated with antibody responses (**Figure S5D**). Altogether, our data points to a fine-tuned system by which the diversity of the microbiome composition differentially influences the outcome of vaccine-induced antigenspecific IgG titers: e.g. reduced antigen-specific IgG titers when the microbiome is disrupted by antibiotic treatment, but restoration of optimal responses when the vaccine is administered after the microbiome is allowed sufficient time to recover.

### Vancomycin Disrupts Vaccine Outcome In Rhesus Macaques

In addition to studying mouse models, we further evaluated whether our observations of vancomycin-induced microbiome disruption and subsequent vaccine hyporesponse were replicated in an NHP model (i.e. rhesus macaque). Rhesus macaques are an important translational animal model with a human-like immune system and microbiome composition. We orally treated groups of 4 vaccine-naïve animals with vancomycin or a vehicle control for 14 days, followed by 1 day of no treatment, prior to each of three doses of intramuscular immunization with the clinically relevant quadrivalent influenza vaccine (Fluzone 2018-2019 season; chosen for its clinical relevance over ovalbumin), given on Days 0, 28, and 84 (**Figure 3A**). While we observed a robust increase in influenza-specific IgG titers in vehicle-treated control animals (peaking at 7 days after the second and third doses), the vancomycin-treated animals had a reduced antibody response (**Figure 3A**). To validate the effect of vancomycin treatment on the microbiome, we longitudinally evaluated the gut microbiome composition by 16s rRNA gene sequencing of stool samples from all animals. As expected, after each 14-day vancomycin treatment regimen (Days 0, 28, 84), the microbial richness was reduced approximately 2-fold compared to vehicle-treated animals or other timepoints (**Figure 3B**) and the microbial communities clustered separately from all other samples (**Figure 3C**).

**Figure 3.**
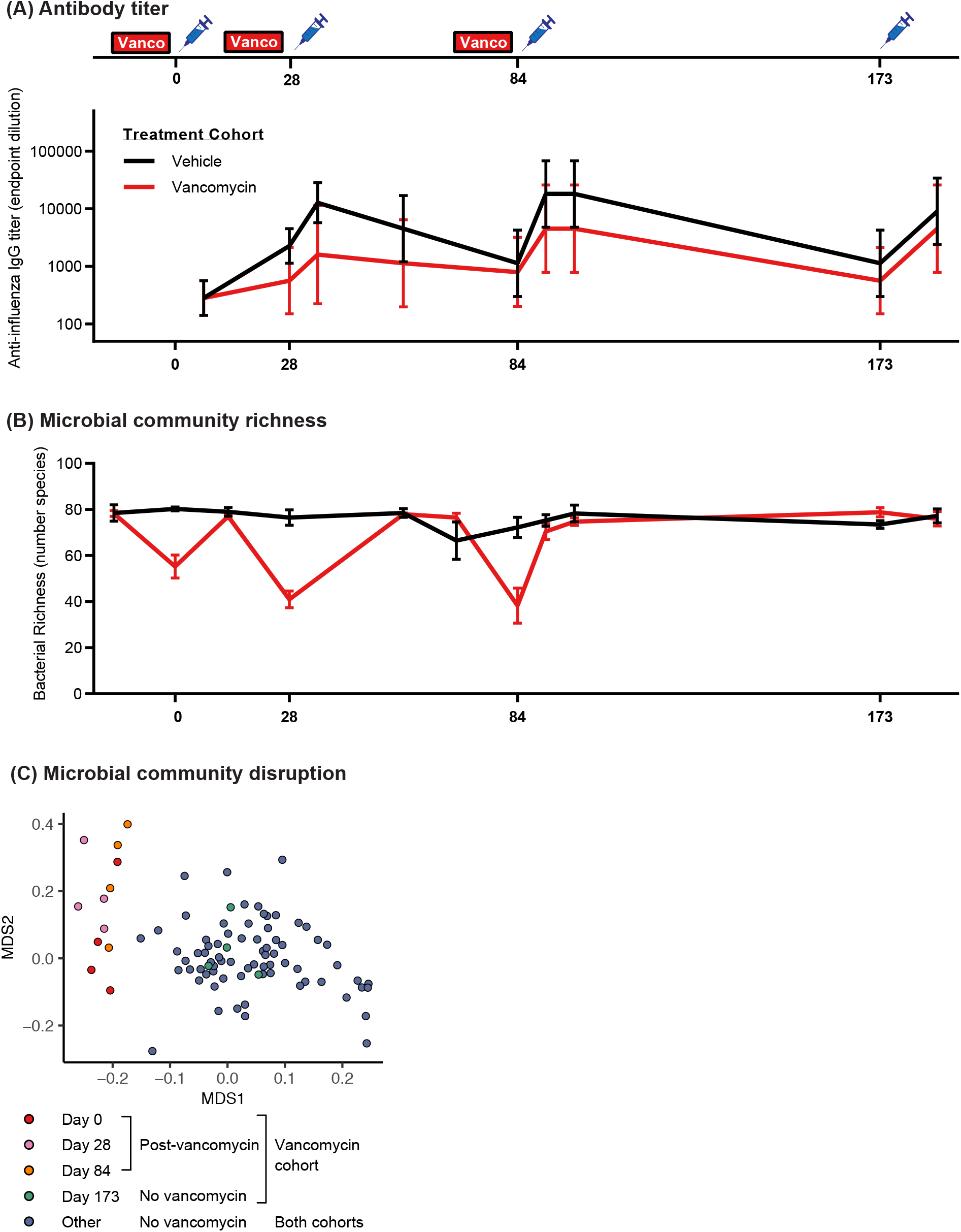
Vancomycin treatment before vaccination reduces influenza-specific antibody titers. (A) Rhesus macaques were orally treated with three 14 day courses of vancomycin (red rectangles), followed by one day without vancomycin, before vaccination with quadrivalent influenza vaccine (blue syringes). A fourth vaccination was performed without vancomycin pre-treatment (Day 173). Serum anti-influenza IgG was measured at multiple timepoints. Data are geometric mean and standard deviation. (B) Fecal samples collected throughout the study were subjected to 16s rRNA gene sequencing, and OTU richness (alpha diversity) was plotted. Data are mean and standard error. (C) Non-metric multidimensional scaling ordination of macaque microbiome communities, with community dissimilarity measured using the Bray-Curtis algorithm (beta-diversity). Samples from the vancomycin cohort taken at the time of vaccination after vancomycin treatment (Days 0, 28, 84) are shown in warm colors, while those taken at vaccination without preceding vancomycin treatment (Day 173) are shown in green. Additional samples from the vancomycin cohort between vaccinations, as well as all samples from the vehicle cohort, are shown in blue and labeled in the key as “Other”.

Because in mice we found that microbiome recovery before vaccination prevents hyporesponse, we administered a fourth booster dose without vancomycin pre-treatment on Day 173 to investigate whether microbiome recovery in macaques previously treated with vancomycin was sufficient to restore vaccine responsiveness. In contrast to the previous vaccination timepoints, the microbiome was similar in both cohorts (**Figure 3**). Strikingly, the vancomycin cohort produced influenza vaccine-specific IgG titers approaching the levels of the NHPs in the vehicle cohort. This crucial observation suggests that microbial dysbiosis at the time of vaccination disrupts vaccine response, but that recovery of the microbiome after antibiotic treatment prior to immunization may lead to successful vaccination responses.

### Vancomycin Impacts Key Innate Immune Pathways in the Mesenteric Lymph Nodes

Together our data suggested that vaccine response is associated with the state of the microbiome at the time of immunization. While measurement of antigen-specific IgG titers served as a surrogate for immune tone in our studies, the specific immune pathways altered upon antibiotic treatment in systemic immune cells or the gastrointestinal mucosal compartment remained unclear. To this end, we treated mice with either vancomycin or a water control for seven days, followed by one day of antibiotic recovery before harvesting the small intestines, mesenteric lymph nodes, spleens, and whole blood. RNA sequencing analysis revealed changes in host gene expression in the small intestine (8 genes), spleen (11 genes), blood (8 genes), and mesenteric lymph nodes (367 genes) due to the perturbation of the gut microbiome.

Within the mouse mesenteric lymph node dataset, we identified 753 Gene Ontology (GO) biological processes that were enriched using gene sent enrichment analysis implemented in ToppGene (44). Clustering of GO terms revealed that vancomycin mediated perturbation of the gut microbiome led to changes in modulated metabolism, signaling, trafficking, differentiation, and cell death pathways, as well as several immune-related pathways (**Figure S6**). Sub-clustering of theimmune-related GO terms highlighted the following immune functions: cytokine response and production, response to bacteria, T-cell adhesion and activation, reactive oxygen and nitrogen species, and inflammatory signaling (**Figure 4A**).

**Figure 4.**
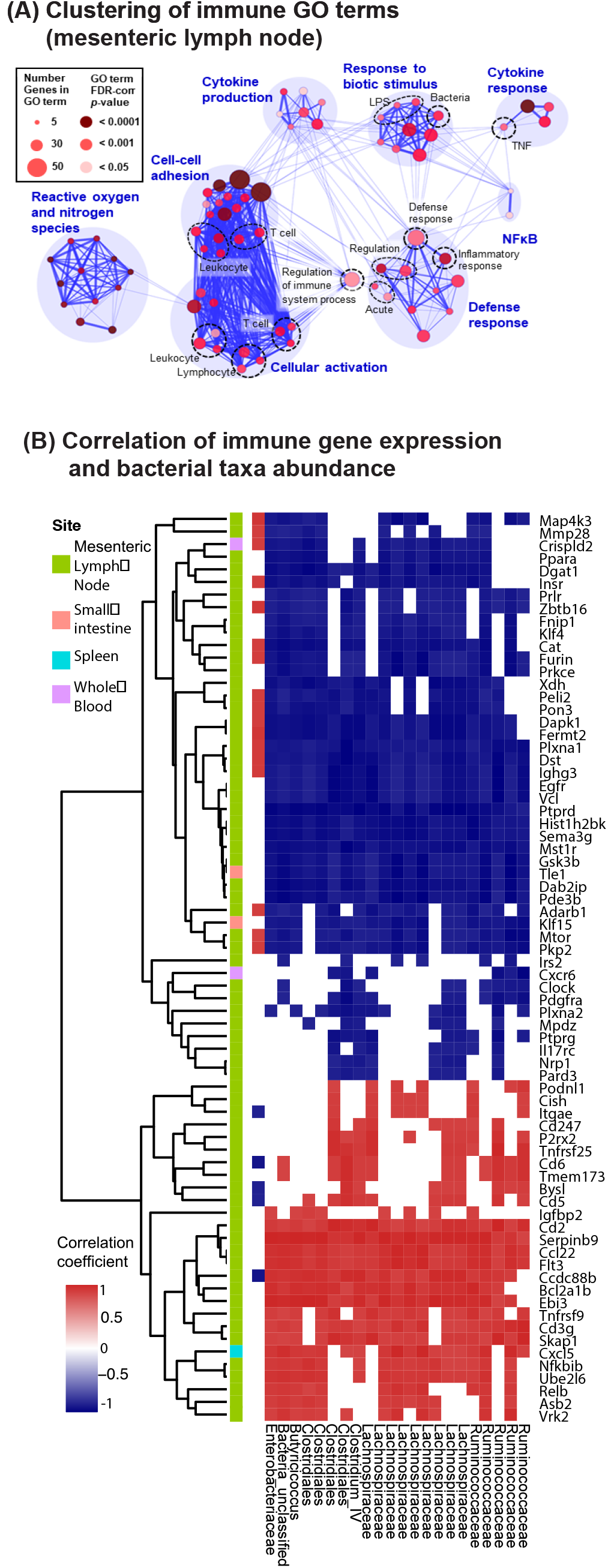
Expression of innate immune response genes is associated with differential abundance of particular bacterial taxa. BALB/c mice were orally treated with vancomycin for 7 days, followed by 1 day without antibiotic treatment before feces and tissues were harvested for bacterial 16S rRNA gene sequencing and host RNA-sequencing. (A) Visualization of immune pathways altered by vancomycin treatment in the mesenteric lymph node. Gene set enrichment analysis was used to identify GO terms enriched in the mesenteric lymph nodes RNA-sequencing dataset (see Figure S6), and enriched immune-related GO terms were clustered based on included differentially expressed genes. Node size and color respectively denote the number of genes present and the statistical significance of enrichment for that GO term; thickness of blue lines represents the similarity between nodes. (B) Differentially expressed genes with immune-related GO terms from the mesenteric lymph node (n=130), as well as immune-related genes differentially expressed in either the small intestine, spleen, or whole blood (n=7) were selected for analysis of association with bacterial OTU (operational taxonomic unit) differential abundance. The Spearman correlation coefficient between abundance of indicated taxa and gene expression across vancomycin- or mock-treated are shown for positive (red) and negative (blue) correlations. Only pairs that are statistically significant (p < 0.01) and with correlation coefficient values beyond 0.8 or −0.8 are shown. Only OTUs correlated with more than 50 genes, or genes correlated with more than 5 genes, are shown. Genes were hierarchically clustered based on correlation with OTUs.

Although our previous analyses did not identify individual bacterial taxa correlated with antibody titer after vaccination, we hypothesized that some taxa could be associated with individual immune pathways. We therefore examined whether vancomycin-induced changes in immune-related gene expression were correlated with the abundance of particular bacterial taxa. We found that specific OTUs within *Lachnospiraceae, Ruminococcus, Clostridia*, and *Enterobacteriaceae* were associated with genes involved in innate and adaptive cell polarization and cytokine/chemokine production. More specifically, we found that several genes associated with productive activation of the immune system were decreased [Asb2 (45), Cxcl5 (46), Tnfrsf9 (47), Bcl2a1b (48), Ccdc88b (49), Flt3 (50)], while genes associated with dampening the immune response were upregulated [Mmp28 (51, 52), Crispld2 (53)]. (**Figure 4B**). This important finding implicates particular bacterial taxa as key immunomodulatory agents and suggests these taxa may work in concert to set the immune “tone” for subsequent immune responses including vaccination. The correlation of multiple bacteria with multiple diverse immune functions also highlights the complexity of microbially-modulated immunity, and the need to continue exploring these underlying dynamics at the systems biology level.

## Discussion

Interactions between the gut microbiota and host play a fundamental role in local and systemic immunity and homeostasis (54–56). Recent reports have employed antibiotics to link microbiome perturbation and the host response to viral infections (57, 58), immune checkpoint inhibitor therapy (59–61), and autoimmune/allergic diseases (39, 42). Several groups have already demonstrated an effect of antibiotic-mediated microbiome perturbation on response to vaccination (22, 24, 27–29, 62, 63), although specific mechanisms and dynamics underlying the vaccine-modulatory effects of antibiotics remain an area of active investigation.

Here we report that differentially perturbing the microbiome, either by using increasingly disruptive antibiotics, decreasing the interval between vancomycin treatment and immunization, or by increasing the duration of vancomycin treatment, proportionally decreased the response to vaccination. Conversely, recovery of the microbiome allowed for recovery of vaccine response. We did not observe relevant associations between vaccine response and specific taxa or microbial metagenomic pathways, which supports the idea that multiple, possibly redundant, microbial factors may modulate vaccine response, rather than specific microbes, in agreement with other preclinical studies of vaccine response (22), tumorigenesis (64) and pathogenesis (65). Indeed, clinical studies linking the microbiome and vaccine response do not reveal a consistent link to a specific microbial signature supporting the existence of redundant overlapping species and pathways.

While previous publications in the microbiome-vaccine research field have employed chronic treatment of combination antibiotics (22–24, 27, 28, 38, 62, 63), we took a broader approach in which we demonstrated individual antibiotics have differential levels of impact on two different strains of mice (BALB/c and C57BL/6). We also note that while the majority of research in this space has focused on aluminum-based adjuvants and inactivated/attenuated viral vaccines (68), we included subunit vaccines adjuvanted with aluminum or novel adjuvants like lipid nanoparticles, as well as inactivated influenza vaccine. Furthermore, we demonstrate vancomycin’s effect on both the model subunit antigen ovalbumin and the clinically relevant hepatitis B surface antigen. Additionally, although the majority of our studies measured IgG titers as a proxy for vaccine response, we also evaluated antigen-specific T cell responses and found appreciable decreases in vancomycin-treated mice. Thus, our data robustly support the conclusion that antibiotic treatment disrupts vaccine response across a variety of formulation measurements and animal models.

Our data support the hypothesis that microbial communities affect immune outcomes by modulating the underlying immune tone, or state of responsiveness. Vancomycin depletion of the gut microbiome diversity modulated expression of key host genes which likely contributed to the immune tone and inflammatory response, which could lead to subsequent vaccine hyporesponsiveness. We also found correlations between altered immune gene expression signatures and certain bacterial taxa. This observation suggests diverse bacteria may work in concert to affect immune tone, which could explain the lack of consistent microbial signatures associated with vaccine response (67). Although most studies in the microbiome-vaccine field rely on 16S rRNA sequencing to identify perturbations in bacterial communities, we additionally performed shot-gun metagenomic sequencing to reveal potential pathways in the microbiome that correlate with antigen-specific IgG titers. This method provides a unique advantage to identify microbial pathways that are potentially indicative of optimal health across broad measures of immune tone. To further dissect the role(s) of the heterogeneous microbiota during response to vaccines, it will be critical to embrace functional and systems biology approaches.

Emerging technologies will play a key role in future investigations of microbiome dynamics and vaccine response. While our approach here using antibiotic treatment facilitated generation of distinct and diverse microbial communities and revealed a spectrum of effects on vaccine response, our current data cannot completely rule out direct effects of antibiotic treatment on the immune system (60, 72, 73). An antibiotic-free model, such as fecal transplant of microbiota into germ-free mice, has been used to demonstrate microbiome-dependent changes in vaccine response and will also be an important next step in our current research program. In addition, observations derived from mouse models must be more extended to human clinical studies (2) to verify that conclusions are relevant for human health.

A unique component of this work is the use of rhesus macaques, an important preclinical model for human vaccines. We found that vancomycin treatment negatively impacted response to inactivated influenza vaccine in rhesus macaques, which mirrors a recent publication that reported a cocktail of antibiotics including vancomycin impacted influenza vaccine outcome in healthy adults with low pre-existing immunity (2). In our study, a booster immunization without prior antibiotic treatment appeared to improve antibody titers. Together, these data highlight the importance of understanding the temporal dynamics of antibiotic treatment and the microbial community when administering vaccines.

Altogether, we report evidence that antibiotic-mediated microbiome perturbation can greatly impact vaccine outcome, but that recovery of the microbiome after cessation of treatment may prevent antibiotic-induced vaccine hyporesponse in adult animals; it remains unclear whether or not this is the case in neonates (23). In addition, our data uncovers important host pathways and immune mediators that may be critical for optimal vaccine outcome. Our results provide support for the importance of understanding antibiotic usage and the microbiome in the context of vaccine discovery. Our results predict that further investigations may identify novel host and microbial factors that are critical for host immunity, and result in the development of more effective vaccines or vaccine co-therapies to address global unmet needs.

## Materials & Methods

### Mice

All animal work was approved by the Institutional Animal Care and Use Committee (IACUC) of Merck & Co., Inc., Kenilworth, NJ, USA in accordance with the recommendations in the Guide for the Care and Use of Laboratory Animals. Specific pathogen-free (SPF) female C57BL/6 and BALB/c mice, aged 5-9 weeks, were purchased from Taconic Biosciences (Rensselaer, NY, USA). Mice were acclimated for 1-2 weeks before experimental treatments.

### Rhesus Macaques

Animal studies involving rhesus macaques were performed at the New Iberia Primate Research Center (NIRC-New Iberia, LA) in accordance with relevant guidelines using protocols approved by NIRC and the Institutional Animal Care and Use Committee of Merck & Co., Inc., Kenilworth, NJ, USA. Female Indian rhesus macaques were weight and age matched for each group. All animals were influenza naïve prior to study start. For each antibiotic course, vancomycin (15 mg/kg) was delivered orally once daily for 14 days. Fluzone quadrivalent influenza vaccine (Sanofi Pasteur; 2018-2019 season) was administered as a single injection per dose, intramuscularly on the right deltoid, in a total volume of 500 μl. Blood for serum and stool were collected at various timepoints and frozen at −80 °C.

### Antibiotic Treatments

Mice were treated with a single antibiotic in drinking water for 1-7 days as indicated in the figure legends. The following antibiotics were used: ampicillin [1 g/L]; clindamycin [1 g/L]; enrofloxacin/Baytril [0.27 g/L]; metronidazole [1 g/L]; neomycin [1 g/L]; nystatin [0. 5 g/L]; tobramycin [1 g/L]; vancomycin [0.5g/L]. Antibiotic-containing drinking water was replaced every 2-3 days. All antibiotics were of pharmaceutical grade and were purchased from United States Pharmacopeia (USP) or Sigma-Aldrich. Prior to vaccination, mice were supplied drinking water without antibiotic for 1-21 days, as indicated in the figure legends.

### Immunizations, Vaccine Antigens & Adjuvants

Mice were immunized by IM injection of 50 μL into each hind limb quadriceps (100 μL total dose per animal per immunization). The following antigens and adjuvants were resuspended in endotoxin-free PBS (Life Technologies) and used for vaccination studies as indicated in figure legends: endotoxin-free ovalbumin (OVA) from Hyglos GmbH [10 μg / 100 μl dose]; recombinant hepatitis B surface antigen (HBsAg) manufactured by Merck & Co., Inc. in West Point, PA, USA [0.2 μg / 100 μl dose]; Imject© Alum from ThermoFisher according to manufacturer’s specifications; lipid nanoparticles based adjuvant VA-510 (LNPs) prepared at Merck & Co., Inc. in West Point, PA, USA as described (76–78) [125 μg / 100 μl dose].

### Collection of Stool and Tissues

Whole blood was collected and mixed with sodium citrate. For serum, blood was collected and allowed to clot. For mice, fresh stool pellets were collected as they were excreted and flash frozen without any buffers/stabilizers. For rhesus macaques, stool samples were collected using manual evacuation method and frozen immediately without any buffers/stabilizers. Spleens, mesenteric lymph nodes, and small intestines without contents, were excised and flash-frozen in liquid nitrogen. All tissues were stored at −80 °C until used for analysis.

### Antigen-Specific Antibody Titers

Sera was collected at the timepoints indicated in the figures and text above. Antigen-specific antibody titers were measured in duplicate via endpoint titer enzyme-linked immunosorbent assay (ELISA), following the Mouse Anti-OVA IgG Antibody Assay Kit (Chondrex), or as previously described (76).

### Pharmacokinetics

Tissues were homogenized in 3-9 volumes DPS buffer (w/v) and extracted by protein precipitation using 3 volumes methanol (v/v) containing a generic internal standard mixture. Extracted samples were quantified by LC-MS/MS on a Waters Acquity UPLC system (Milford, MA, USA) with mobile phase A consisting of 0.1% formic acid in water, and mobile phase B consisting of 0.1% formic acid in acetonitrile using a Waters HSS T3 (100 Å, 1.8 μm, 2.1 mm X 50 mm) column maintained at 30 °C with gradient elution. The analyte and internal standards were detected via tandem mass spectrometry employing an AB/Sciex 5000+^™^ triple quadrupole system (Concord, ON, Canada) using Multiple Reaction Monitoring (MRM).

### T-cell Activity Quantification

Spleens from vaccinated mice were collected at the indicated timepoint, mechanically disrupted, and treated with ACK (Ammonium-Chloride-Potassium) buffer to lyse erythrocytes. Splenocytes were cultured at 37 °C, 5% CO_2_ in RPMI media with 10% FBS and 1.25 μg/ml each α-CD28 (clone 37.51, BD) and α-CD49d antibodies (clone R1-2, BD). MHCI-specific ovalbumin SIINFEKL/OVA_257-264_ or MHCII-specific ovalbumin peptide OVA_323-339_ were added at 2 μg/ml (Invivogen). Phorbol 12-myristate 13-acetate (Promega) and ionomycin (Tocris) were added at 1.25 μg/ml and 0.02 μg/ml, respectively. After 24 hours, interferon-gamma (IFN-γ) in the supernatant was quantified using the mouse IFN gamma ELISA (AbCam).

### DNA extraction and 16S rRNA Gene Sequencing

DNA from 1-2 stool pellets was extracted using the QIAamp DNA Stool Mini Kit (Qiagen) as per the manufacturer’s instructions. The V4 region of the 16S rRNA gene of the stool samples was amplified on the Illumina MiSeq platform using their standard 2×250bp paired-end protocol and the 515F and 806R 16S rRNA gene sequence primers (79). Resulting DNA sequences were processed as previously described (80). Briefly, we used Mothur (v1.39.5) to assemble the paired-end reads (requiring complete overlap) and perform quality filtering. Sequences were aligned to the SILVA 16S rRNA gene sequence database (v132) and filtered to remove chimeras and lineages of archaea, chloroplasts, and mitochondria. OTUs were clustered at 97% similarity using the OptiClust algorithm, and reads were aligned to OTUs for quantification. OTU tables and taxonomic assignments were exported for downstream analysis in R.

The rhesus macaque stool sample 16S rRNA gene sequenced were bioinformatically processed using the standard CosmosID protocol. Briefly, OTUs were identified using a closed-reference approach and 97% sequence similarity using Qiime (81).

### Shotgun Metagenomic Sequencing and KEGG pathway enrichment analysis

Reads were randomly amplified for sequencing using the Illumina TruSeq library kit. The paired-end sequencing was performed on the Illumina HiSeq platform using a universal primer in one direction, and a set of standard index primers in the reverse direction. Sequences were processed for quality using bbduk for adapter removal and quality trimming (ktrim=1, mink=17, maq=30, minlen=75), and fastqc for general quality visualization. Bowtie2 was used to remove reads that aligned with similarity to the PhiX and human genome. Details can be found in the supplemental analysis code.

Metagenomic sequences were aligned to the KEGG database through the public API (accessed August 12, 2020) and using the diamond alignment algorithm (82–85). KEGG orthology (KO) counts were generated by compressing gene counts and equally splitting counts when genes were present in multiple KO groups, or vice versa. KO counts were normalized using the MUSICC algorithm (86). KEGG pathway enrichment was calculated using the Generally Applicable Geneset/Pathway (GAGE) analysis package, available through R Bioconductor. The enrichment calculation for each antibiotic treatment group was performed in reference to the water control group, using the “unpaired” comparison and “same direction” parameters (87).

### RNA Extraction, Sequencing, and differential gene expression analysis

Total RNA was extracted from spleens, mesenteric lymph nodes, and small intestine without contents using the RNeasy Mini Kit (Qiagen) as per manufacturer’s instructions. Whole-blood was collected using RNAprotect Animal Blood Tubes (Qiagen) and processed as per manufacturer’s instructions. RNA quality and concentration was measured using an Agilent 4200 TapeStation (Agilent Technologies) as per manufacturer’s instructions and an RNA integrity number (RIN) cutoff of 6 was used to progress samples to sequencing. RNA was sequenced by BGI Genomics (Shenzhen, China). FASTQ files were trimmed and mapped with OmicSoft (v10.1) [OmicSoft Corporation]. Trimming was performed with a 5bp window starting from the 3’ end. Using this sliding window, if 2 or more bases in this 5bp window were below a PHRED quality score of 15, those bases and any base pair to the right were removed.

For analysis of differential gene expression, genes were included if the read count was above 1 read count per million in at least 5 samples of the group. Differential expression was calculated using EdgeR (v3.20.9) which leverages Limma dependencies (v3.34.9) in the R statistical computing environment [R version 3.4.3 (88)]. EdgeR moderates the degree of overdispersion using an overdispersed Poisson model of biological and technical variability coupled with an empirical Bayes method. Library size normalization factors were generated using the trimmed mean of M-values (TMM) method (trimmed mean of M-values) from the “calcNormFactors” function in the edgeR library and used as covariates in modeling. Data was then fit to a generalized linear model (GLM) and a GLM likelihood ratio test was performed to determine if the coefficient representing the contrast between the conditions of interest was equal to 0 indicating no differential expression. Significance for differential expression was set at an FDR corrected p-value < 0.05.

### GO term analysis & clustering

Gene Ontology (GO) terms were analyzed for enrichment using the ToppFun program from ToppGene (44). Differentially expressed genes (FDR corrected p-values < 0.05) were provided to this program to assess enrichment in the following GO categories: Biological Process, Molecular Function, and Cell Component. A term was considered significantly enriched with FDR (Benjamini and Hochberg) corrected p-value < 0.05. The mesenteric lymph node GO term network was constructed using the Cytoscape application EnrichmentMap (89), using FDR (BH) cutoff of 0.035 and overall edge cutoff of 0.8, and clusters were identified using ClusterMaker— MCL Cluster (90) with granularity parameter of 1.5. Clusters were manually annotated. A new network was constructed from GO terms in immune-related clusters using EnrichmentMap with an overall edge cutoff of 0.6, and clusters were manually identified and annotated.

### Statistical Analyses

Statistical analyses were performed in the R statistical computing environment (91). The statistical tests, multiple hypotheses corrections, and other details are identified above, in association with the results, and the details can be found in the associated analysis code.

## Data & Code Availability

All sequencing data is available through the Sequence Read Archive at PRJNA718580. Code is available through GitHub at https://github.com/Merck/AbVax_Manuscript.

## Author Contributions

Contributions assigned using the CRediT Contributor Role Taxonomy system. G Swaminathan, M Citron, JE Norton, JM Maritz, CH White, S Kommineni, X Liang, DA Gutierrez, O Danilchanka, CH Woelk, DJ Hazuda, and GD Hannigan contributed to **study conceptualization**. **Data was analyzed, curated, and managed** by G Swaminathan, M Citron, J Xiao, JE Norton, AL Reens, BD Topçuoğlu, JM Maritz, KJ Lee, DC Freed, TM Weber, CH White, M Kadam, and GD Hannigan. G Swaminathan, M Citron, J Xiao, JE Norton, JM Maritz, KJ Lee, DC Freed, TM Weber, CH White, S Kommineni, X Liang, and GD Hannigan **performed the investigation**. G Swaminathan, M Citron, J Xiao, JE Norton, BD Topçuoğlu, JM Maritz, KJ Lee, DC Freed, TM Weber, CH White, S Kommineni, X Liang, DA Gutierrez, GD Hannigan, E Spofford, E Bryant-Hall, G Salituro, O Danilchanka, and J Fontenot **developed and designed methodologies**. **Project administration** was managed by G Swaminathan, M Citron, BD Topçuoğlu, JM Maritz, GD Hannigan, E Spofford, and E Bryant-Hall. **Resource were provisioned** by J Xiao, JE Norton, BD Topçuoğlu, JM Maritz, DA Gutierrez, CH Woelk, GD Hannigan, E Spofford, E Bryant-Hall, Gino Salituro, O Danilchanka, and J Fonenot. **Software implementation and support** were provided by AL Reens, BD Topçuoğlu, JMM Maritz, and GD Hannigan. **Project was supervised** by G Swaminathan, M Citron, DA Gutierrez, DJ Hazuda, CH Woelk, and GD Hannigan. **Validation and verification** was done by G Swaminathan, J Xiao, JE Norton, AL Reens, BD Topçuoğlu, JM Maritz, J Fontenot, and GD Hannigan. **Data visualization and presentation were organized** by AL Reens, BD Topçuoğlu, JM Maritz, GD Hannigan. **The original manuscript draft was prepared** by G Swaminathan, AL Reens, and GD Hannigan. **Manuscript review and editing** were performed by G Swaminathan, AL Reens, BD Topçuoğlu, CH Woelk, and GD Hannigan.

## Acknowledgements

The authors would like to acknowledge members of the laboratories animal research (LAR) division at MRL, West Point & Boston for assistance with rodent experiments. We sincerely thank members of the New Iberia Research Center (NIRC), Louisiana, for assistance with the rhesus macaque studies. We appreciate contributions from Alexander Kulyk, Rebecca Custers-Allen and Jennifer Cho on sequencing related end points. In addition, we would like to thank Sandip Datta, Antonios Aliprantis, Amy Espeseth, Alex Therien, Robin Kaufhold, Julie Skinner, Todd Black, Cameron Douglas, Juliana Malinverni, Jessica Flynn, Lan Zhang, Jill Maxwell, and Andrew Bett for input and thoughtful comments on the study design, interpretation, and/or on the manuscript.

## Supplemental Figures

**Figure S1.**
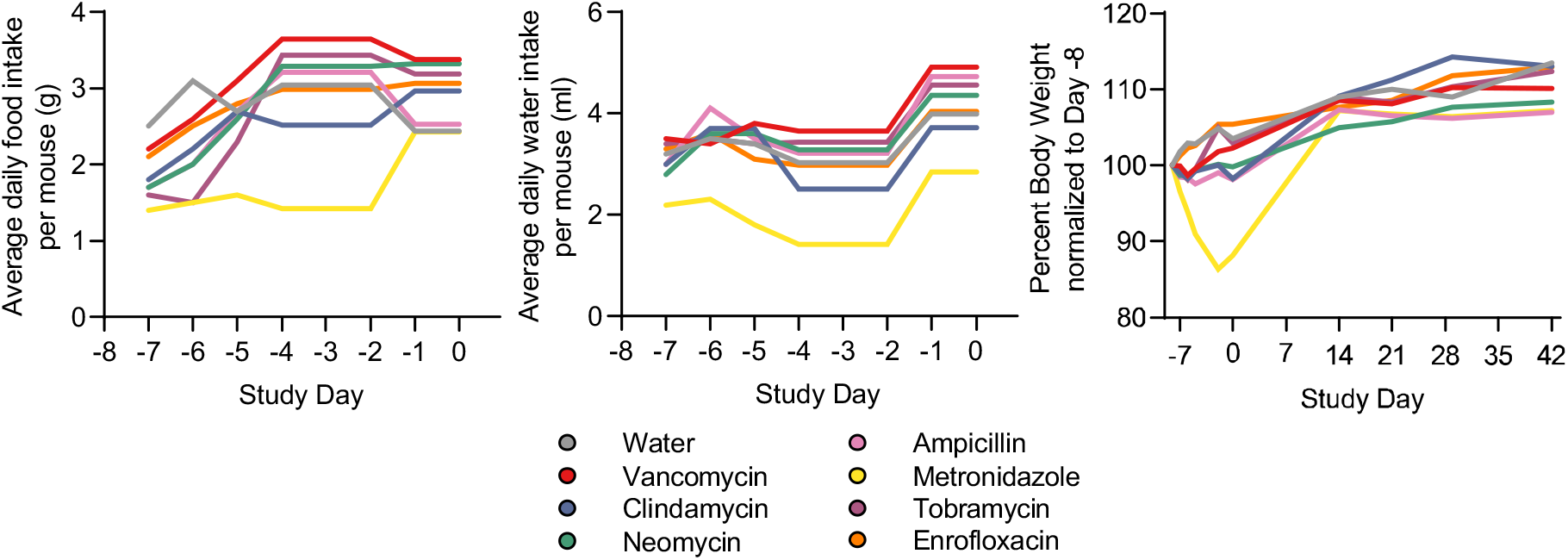
Antibiotic treatment does not affect health measures in BALB/c mice. Food intake, water intake, and body weight normalized to that at the beginning of the study [100%] shown in Figure 1.

**Figure S2.**
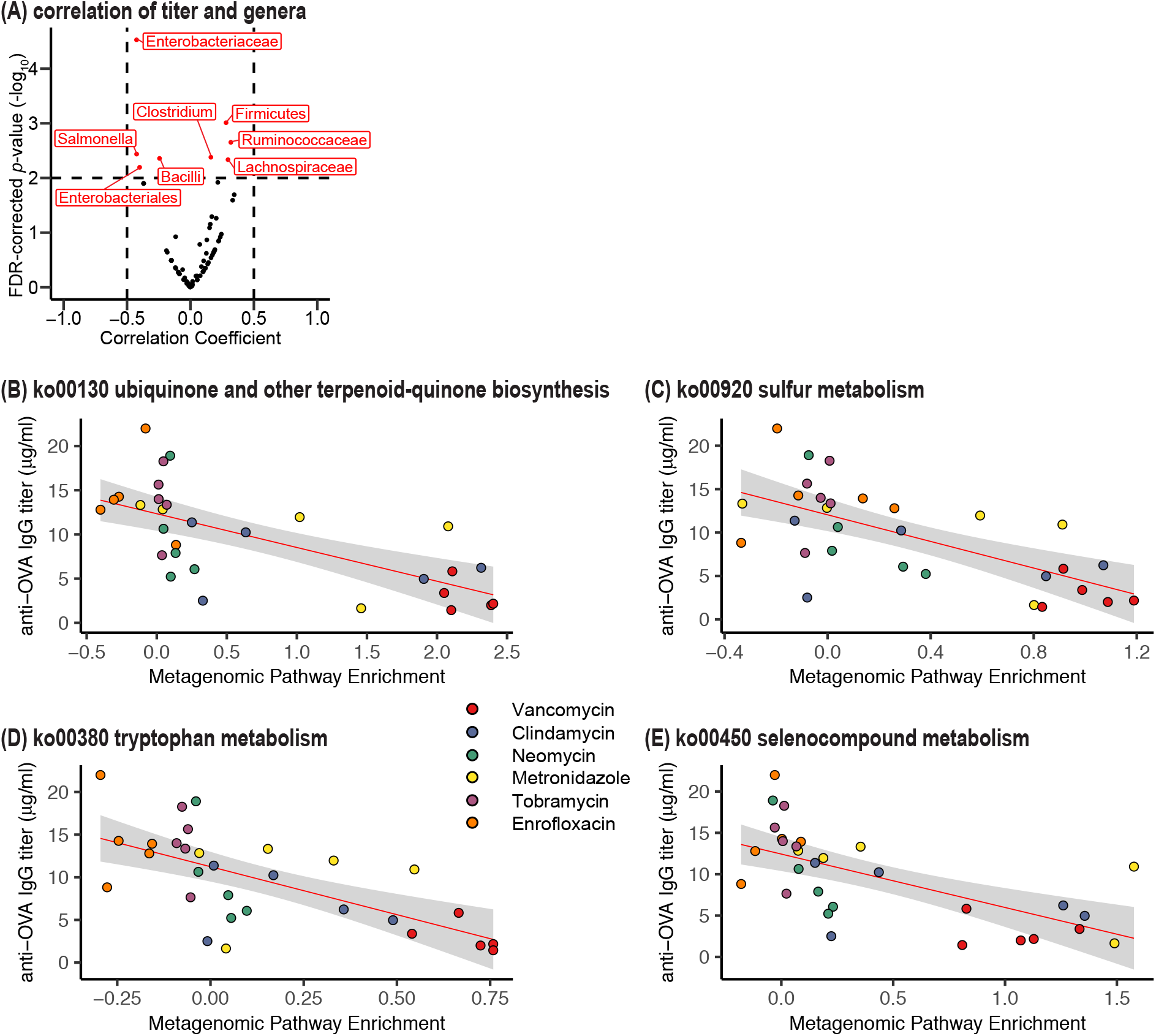
Anti-ova IgG titers do not strongly correlate with abundances of bacterial genera across antibiotic treatments or enrichment of metagenomic pathways. (A) Volcano plot showing correlation analysis of anti-OVA IgG titers and abundances of specific bacterial genera at Day 0. Red dots represent taxa that were statistically significantly correlated (FDR-corrected p-value < 0.01), although the correlation coefficient did not exceed 0.5, indicating negligible relevance. (B-E) Correlation scatter plots of each individual mouse’s OVA-specific IgG titer and metagenomic enrichment of the indicated Kegg orthology pathways relative to water-treated animals. Ampicillin-treated animals were not included in the metagenomic analysis due to poor metagenomic sequence yield. Red line indicates linear regression; gray shading represents 95% confidence interval. Pearson correlation, FDR p-value < 0.005, r < −0.60).

**Figure S3.**
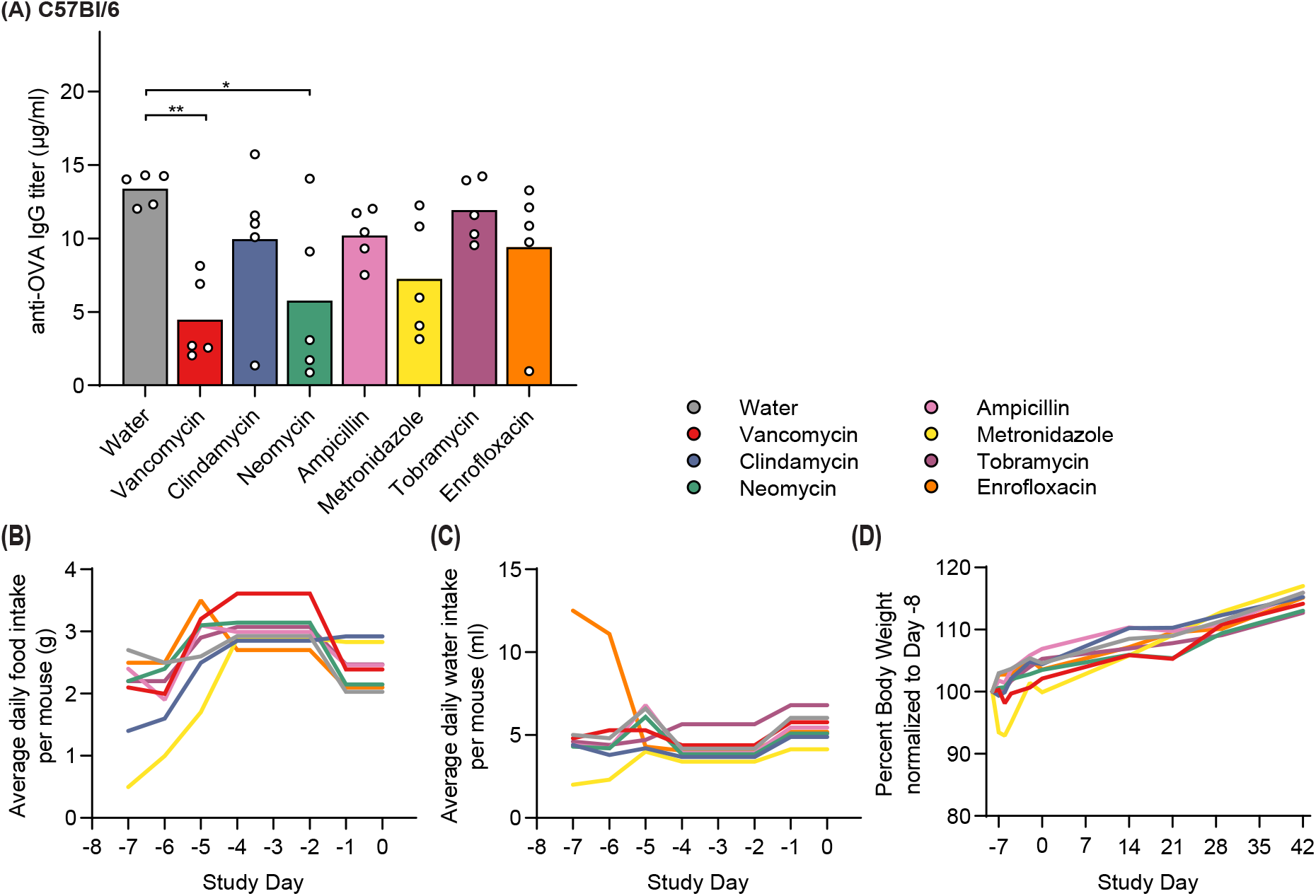
Antibiotic treatment disrupts antibody response to vaccination but does not affect health measures in C57Bl/6 mice. C57Bl/6 mice were treated as described in Figure 1. (A) Serum OVA-specific IgG titers were measured at Day 42 after immunization. Measurements from individual mice (circles) are superimposed on group mean (line). * p < 0.05, ** p < 0.01 compared to water by Dunnett’s test. (B-D) Food intake, water intake, and body weight normalized to that at the beginning of the study [100%].

**Figure S4.**
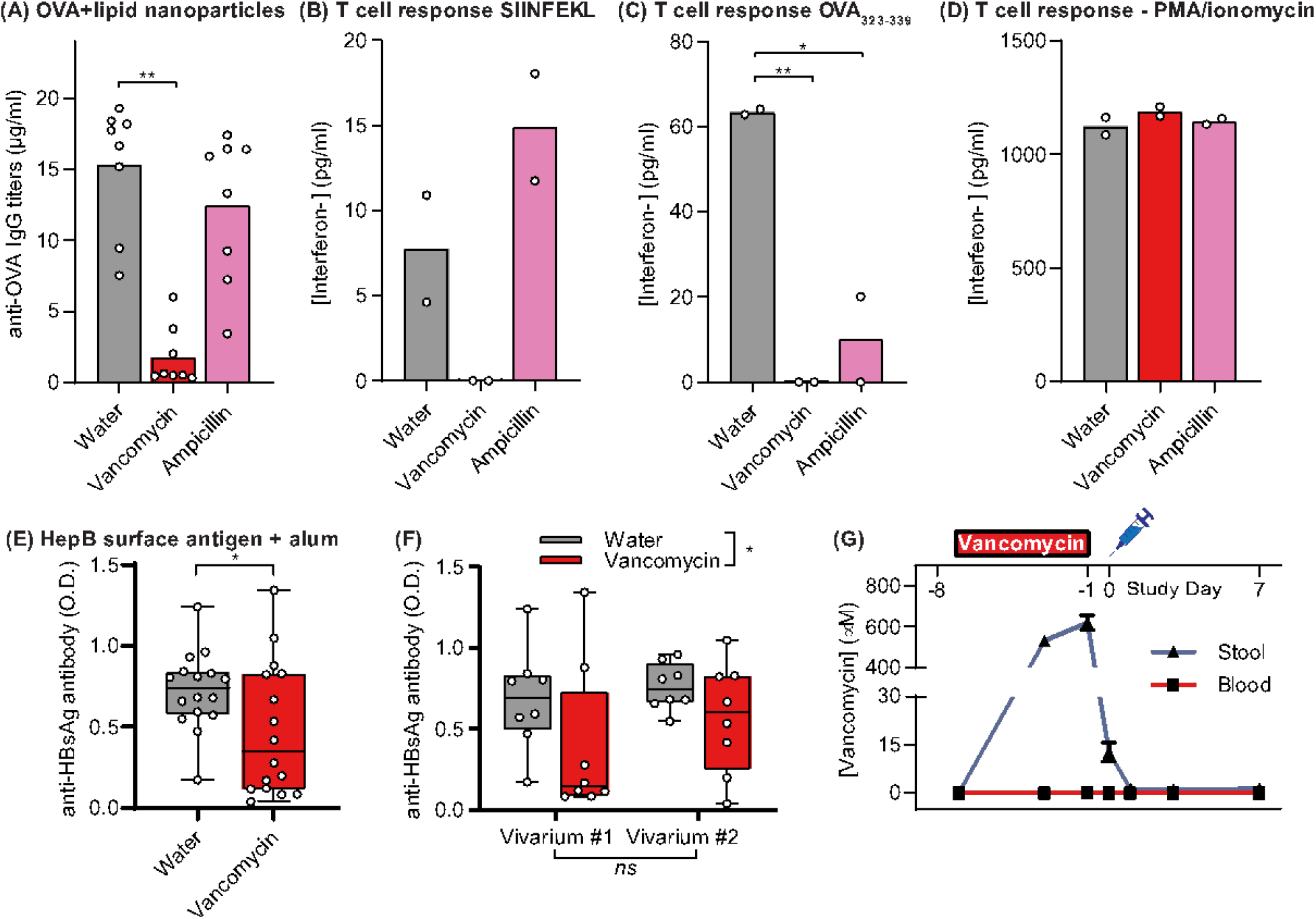
Vancomycin treatment affects response to vaccines with different adjuvants and antigens. (A-D) BALB/c mice were treated with ampicillin or vancomycin for 7 days, followed by one day without antibiotic treatment before immunization with ovalbumin and VA-510 LNPs at Days 0 and 21. (A) Quantification of OVA-specific IgG titers at Day 41. (B-D) Interferon-gamma production by splenocytes from antibiotic-treated mice in response to stimulation with the indicated molecules. (E-G) Separate cohorts of BALB/c mice at vivaria in different states were treated with vancomycin for 7 days, followed by one day without antibiotic treatment before immunization with HBsAg and alum at Days 0 and 21. Serum HBsAg-specific IgG was quantified at Day 35 and plotted across both cohorts (E) or separated by cohorts (F). Measurements from individual mice (circles) are superimposed on group mean (line). Boxplots indicate upper and lower quartiles. (G) Concentrations of vancomycin in mouse stool and blood 7 days before and after the first immunization; data are mean + SEM of 5 mice. * p < 0.05, ** p < 0.01, *** p < 0.001 compared to water by one-way ANOVA with Dunnett’s post-test (A-D), one-sample t test (E), or two-way ANOVA (F).

**Figure S5.**
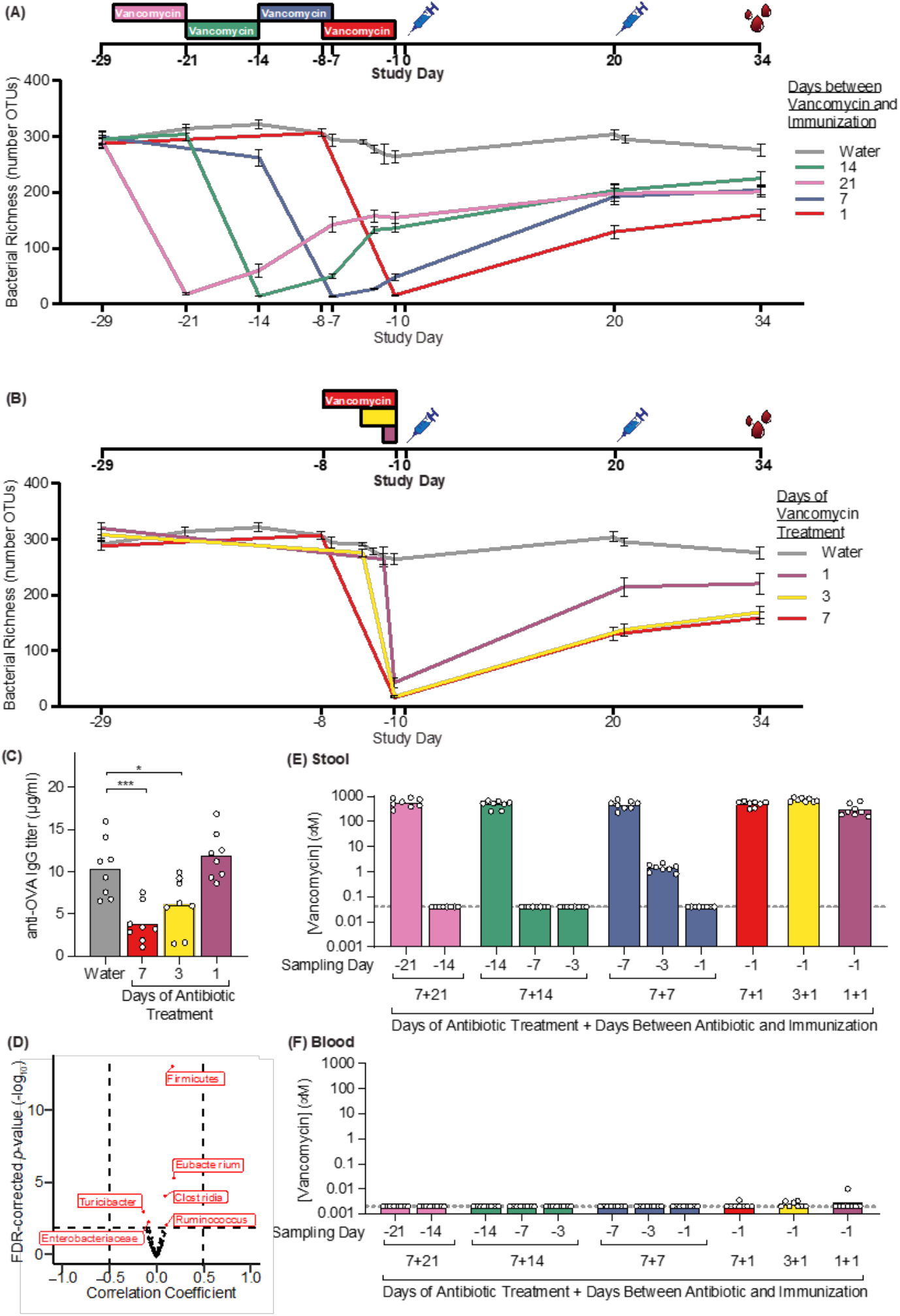
Microbiome recovery after antibiotic treatment. (A) OTU richness (alpha diversity) of groups described in Figure 3. (B,C) In parallel to the samples described in Figure 3, mice were orally treated with vancomycin for 7, 3, or 1 day (rectanges), followed by 1 day without antibiotic treatment, prior to immunization as in Figure 3. Data from the group receiving 7 days vancomycin and 1 day without antibiotic is repeated from Figure 3. (B) OTU richness (alpha diversity) during the study. (C) Serum anti-OVA IgG was measured at Day 34; * p < 0.05, *** p < 0.001 compared to water by Dunnett’s test. (D) Volcano plot showing correlation analysis of anti-OVA IgG titers and abundances of specific bacterial genera at Day -1. Six taxa (red dots) were statistically significantly correlated (FDR-corrected p-value < 0.01; horizontal dashed line), although the correlation coefficient did not exceed 0.5 (vertical dashed lines), indicating negligible relevance. (E,F) Concentrations of vancomycin in mouse stool and blood. Dashed line represents limit of quantification. Measurements from individual mice (circles) are superimposed on group mean (line).

**Figure S6.**
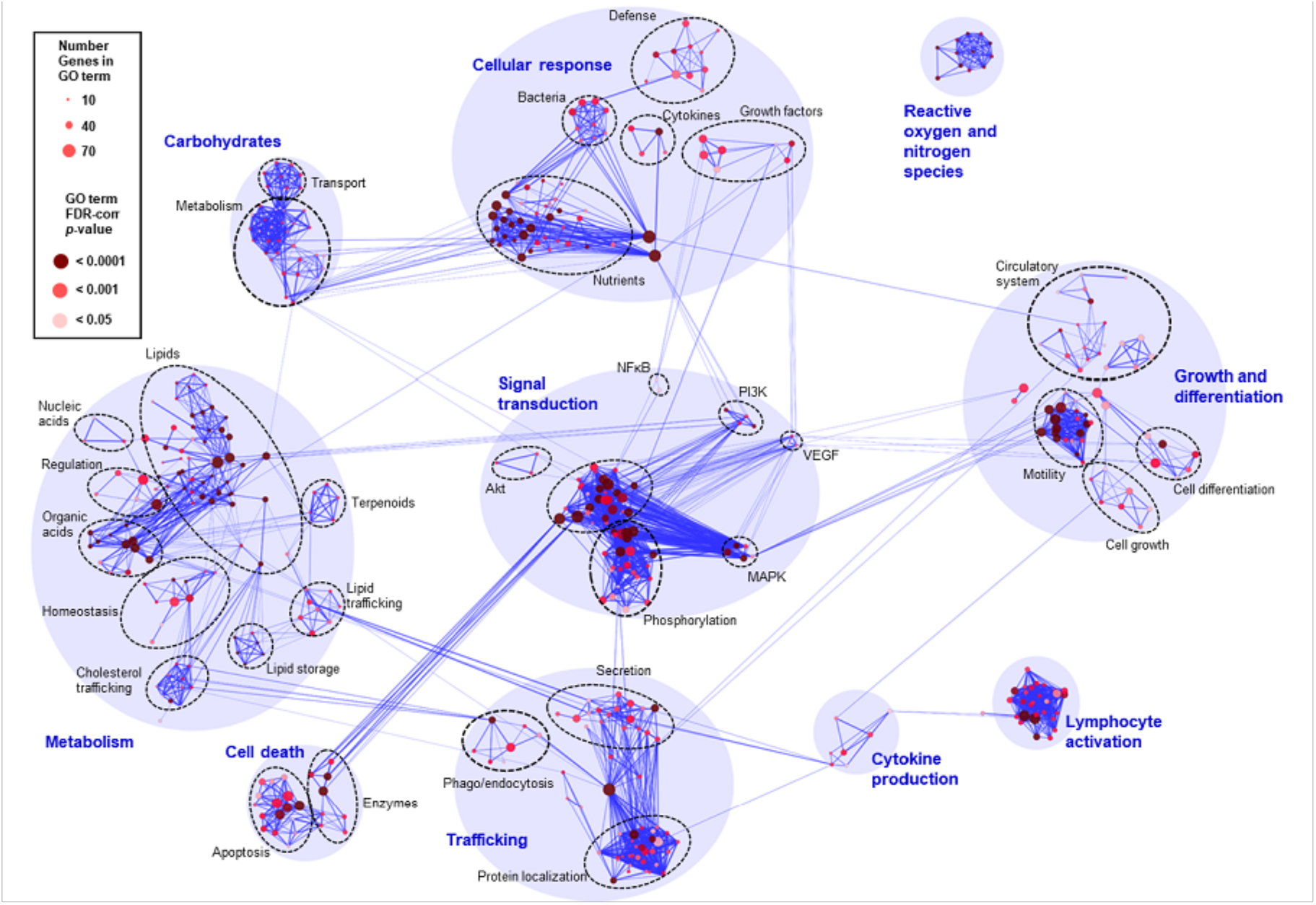
Clustering of enriched GO terms in RNA sequencing of mesenteric lymph nodes from vancomycin-treated mice. Gene set enrichment analysis was used to identify GO terms enriched in the vancomycin-treated mesenteric lymph nodes RNA-sequencing dataset. All enriched GO terms were clustered based on included differentially expressed genes. Node size and color respectively denote the number of genes present in and the statistical significance of enrichment for that GO term; thickness of blue lines represents the similarity between nodes. The following clusters were denoted as “immune-related,” subsampled and clustered (Figure 4A), and genes within were used for correlation analysis with OTU abundances (Figure 4B): Lymphocyte activation, Cytokine production, NFκB (Signal Transduction), Cellular Response, and Reactive oxygen and nitrogen species.

## Supplemental Tables

**Table S1.** Metagenomic KEGG pathways depleted in antibiotics groups, compared to the water control. Pathways shown have FDR corrected p-values < 0.01.

|  | stat.mean | q.val | KOID | treatment |
| --- | --- | --- | --- | --- |
| 1 | -2.377896 | 1.130830e-04 | ko00511 Other glycan degradation | Baytril |
| 2 | -1.833744 | 3.621323e-03 | ko04142 Lysosome | Baytril |
| 3 | -2.461681 | 1.296014e-05 | ko00970 Aminoacyl-tRNA biosynthesis | Clindamycin |
| 4 | -1.924595 | 4.323888e-03 | ko00970 Aminoacyl-tRNA biosynthesis | Metronidazole |
| 5 | -3.116655 | 3.904034e-09 | ko00970 Aminoacyl-tRNA biosynthesis | Neomycin |
| 6 | -2.804875 | 2.016370e-07 | ko00970 Aminoacyl-tRNA biosynthesis | Tobramycin |
| 7 | -2.613916 | 2.015826e-06 | ko00970 Aminoacyl-tRNA biosynthesis | Vancomycin |
| 8 | -1.723265 | 8.585391e-03 | ko04151 PI3K-Akt signaling pathway | Vancomycin |

**Table S2.** Metagenomic KEGG pathways enriched in antibiotics groups, compared to the water control. Pathways shown have FDR corrected p-values < 0.01.

|  | stat.mean | q.val | KOID | treatment |
| --- | --- | --- | --- | --- |
| 1 | 2.583535 | 4.259424e-06 | ko00970 Aminoacyl-tRNA biosynthesis | Baytril |
| 2 | 1.778945 | 9.403102e-03 | ko00040 Pentose and glucuronate interconversions | Clindamycin |
| 3 | 1.956431 | 3.955527e-03 | ko00511 Other glycan degradation | Tobramycin |
| 4 | 2.208632 | 8.705120e-05 | ko02010 ABC transporters | Vancomycin |
| 5 | 2.210136 | 1.030133e-04 | ko00130 Ubiquinone and other terpenoid-quinone biosynthesis | Vancomycin |
| 6 | 1.934026 | 3.585939e-04 | ko02024 Quorum sensing | Vancomycin |
| 7 | 2.016123 | 3.585939e-04 | ko00780 Biotin metabolism | Vancomycin |
| 8 | 1.930946 | 3.585939e-04 | ko00540 Lipopolysaccharide biosynthesis | Vancomycin |
| 9 | 1.897841 | 3.585939e-04 | ko01230 Biosynthesis of amino acids | Vancomycin |
| 10 | 1.908200 | 3.585939e-04 | ko02060 Phosphotransferase system (PTS) | Vancomycin |
| 11 | 1.813439 | 1.029094e-03 | ko00053 Ascorbate and aldarate metabolism | Vancomycin |
| 12 | 1.719883 | 2.587855e-03 | ko04122 Sulfur relay system | Vancomycin |
| 13 | 1.632562 | 3.462776e-03 | ko00790 Folate biosynthesis | Vancomycin |
| 14 | 1.534577 | 6.717780e-03 | ko00400 Phenylalanine, tyrosine and tryptophan biosynthesis | Vancomycin |
| 15 | 1.504306 | 8.055240e-03 | ko05111 Biofilm formation – *Vibrio cholerae* | Vancomycin |

